# Self-Calibrated Hyperspectral Neural Radiance Fields for 3D Reconstruction of Bone and Bone Analogues

**DOI:** 10.64898/2026.06.17.732938

**Authors:** Neetu Sigger, Tuan Thanh Nguyen, Sadaf Ashraf, Gianluca Tozzi

## Abstract

Hyperspectral imaging (HSI) has gained increasing attention for bone assessment because it captures rich wavelength dependent information associated with mineralised tissue. HSI provides detailed spectral information related to material composition, while 3D geometric information supports the analysis of surface morphology and structural detail. However, integrating spectral and geometric information remains challenging, particularly when conventional reconstruction pipelines depend on external pose estimation. To address this challenge, we propose BoNeRF-HS, a self-calibrated hyperspectral neural radiance field for 3D reconstruction. BoNeRF-HS jointly optimises camera intrinsics, volume density, and hyperspectral radiance, removing the need for COLMAP based poses. To improve spectral modelling, we incorporate a gated spectral adapter head that learns wavelength dependent radiance features for hyperspectral view synthesis. We evaluate BoNeRF-HS on a multi-view hyperspectral dataset containing mouse bone, trabecular bone analogue, and cortical bone analogue samples. Experimental results demonstrate that our framework achieves improved reconstruction quality, and better preservation of bone surface details compared with existing approaches.

## Introduction

Recent advances in orthopaedic and trauma surgery have increased the use of intraoperative imaging to provide patient specific structural information during bone related procedures. In particular, fluoroscopy and intraoperative three dimensional cone beam computed tomography (CBCT) are widely used for fracture reduction assessment, implant position verification, and surgical navigation^1–3^. These modalities allow surgeons to assess fracture alignment and identify malreduction or implant malposition before wound closure^4,5^. Although these imaging techniques provide valuable structural information, they are associated with several practical limitations, including ionising radiation exposure, two-dimensional projection ambiguity in fluoroscopic imaging, and limited capability to assess tissue specific spectral responses such as wavelength dependent reflectance, oxygenation, or biochemical composition^1,3,6^.

Optical three dimensional reconstruction has therefore been investigated as a non-contact and non-ionising approach for capturing exposed bone surfaces during surgical procedures^7,8^. Structured light imaging^9^, and multi-view reconstruction^8^ can provide dense surface geometry and may support surface-based registration, surgical navigation, and intraoperative visualisation^7,8^. However, reliable reconstruction of exposed bone remains challenging because bone surfaces often exhibit limited texture, specular reflections, irregular morphology, and non-uniform illumination, which can affect feature matching, depth estimation, and surface reconstruction accuracy^8^. Furthermore, geometric surface models alone do not provide information about tissue composition, spectral response, or surface condition^10,11^.

Hyperspectral imaging (HSI) acquires both spatial and spectral information by capturing a reflectance spectrum at each image pixel^10^. Unlike RGB imaging, which records only three broad colour channels, HSI measures many narrow wavelength bands and can reveal wavelength dependent tissue signatures related to physiology, morphology, and composition^12,13^. This makes HSI particularly attractive for medical and surgical applications, where it may support non-contact and label-free tissue assessment^10,11^. In orthopedic applications, HSI has been explored for several clinically relevant tasks^11,14^. Kistler et al.^14^ used HSI to assess knee cartilage and demonstrated that spectral variations could help differentiate damaged cartilage from healthy cartilage. Barberio et al.^11^ highlighted the role of intraoperative HSI in supporting tissue discrimination during surgery, while Chehade et al.^15^ evaluated lightweight learning models for HS image segmentation and showed that a multilayer perceptron (MLP) could effectively distinguish bone and cartilage regions, demonstrating its potential in musculoskeletal surgical environments.

Despite these advantages, most surgical HSI approaches analyse the scene as a two-dimensional spectral image^10,11,16^. This limits the interpretation of spectral signatures within the three-dimensional structure of the surgical scene. Combining HSI with 3D reconstruction can overcome this limitation by mapping spectral information onto reconstructed bone surfaces, allowing surface geometry and wavelength-dependent tissue responses to be examined within a common spatial representation^17–19^. Previous studies have shown the feasibility of combining surface reconstruction with hyperspectral or multispectral imaging for surgical applications, including endoscopic surface shape reconstruction^17^, light field HSI^18^, and real-time stereo HSI^19^. Nevertheless, 3D hyperspectral reconstruction of exposed bone surfaces remains relatively underexplored, particularly in settings where accurate camera poses are not externally available.

Neural radiance fields (NeRFs) have emerged as a powerful framework for learning 3D scene representations^20^. A NeRF maps the 3D position and viewing direction to radiance and volume density, and uses differentiable volume rendering to synthesise novel views^20^. Recent work has begun to extend neural rendering from RGB images to hyperspectral data^21,22^. Hyperspectral NeRFs that model wavelength dependent radiance and transmittance^23^, while HS-3D-NeRF uses multi-channel NeRFs for joint 3D surface and hyperspectral reconstruction^22^. Recent hyperspectral Gaussian Splatting^24^ methods further explore efficient spectral scene representations for 3D hyperspectral rendering. These methods demonstrate the potential of neural rendering for recovering both geometry and wavelength dependent appearance. However, these hyperspectral 3D methods assume known or externally estimated camera parameters^23,24^. In contrast, self-calibrated RGB NeRF methods such as NeRF–^25^ jointly optimise camera intrinsics, camera poses, and the radiance field without requiring precomputed poses. Building on this idea, our work extends self-calibrated neural rendering to hyperspectral reconstruction.

This paper presents a novel self-calibrated hyperspectral neural rendering framework, called BoNeRF-HS, for 3D reconstruction of hyperspectral images. BoNeRF-HS jointly optimises camera intrinsics, per-view camera poses, volume density, and hyperspectral radiance, removing the need for COLMAP poses. The main stages of the proposed framework are summarised as follows. First, we construct a multi-view hyperspectral bone dataset containing one mouse bone sample and two bone-analogue samples. Secondly, we extend the NeRF–^25^ self-calibrated formulation from RGB to hyperspectral radiance prediction, enabling direct reconstruction of wavelength dependent appearance from HSI. Finally, to better utilise the spectral information across neighbouring bands, a gated spectral adapter head is introduced. This module learns a controlled residual correction to the direct spectral prediction, allowing the model to refine spectral responses while preserving the main hyperspectral radiance estimate. The results show that BoNeRF-HS improves hyperspectral reconstruction quality compared with the other self-calibrated methods, producing sharper rendered views, and better preservation of bone surface details. More broadly, this study demonstrates the potential of self-calibrated hyperspectral neural rendering for non-contact 3D spectral reconstruction of bone surfaces, where geometric structure and spectral responses can be analysed within a unified representation.

## Results

In this section, we present the reconstruction results of the proposed BoNeRF-HS framework and compare them with existing methods using both quantitative metrics and qualitative visualisations.

### Training process

The proposed BoNeRF-HS model was implemented in PyTorch and trained on 120 band HSI. Three specimen types were used in this study, one mouse bone and two bone analogue samples, trabecular bone analogue and the cortical bone analogue. We used approximately 90% of the views for training and the remaining views for testing. The model jointly optimises the hyperspectral radiance field, camera intrinsics, and per-view camera poses. During each training iteration, rays are sampled from the training views, and 128 points are sampled along each ray. Specifically, we use the spectral angle mapper (SAM) loss to preserve spectral-shape similarity between the predicted and ground-truth spectra^26^. The model was trained using the Adam optimiser^27^ for 10,000 epochs. During optimisation, randomly sampled 32 × 32 ray batches were used for each training step.

### Performance evaluation

The performance of BoNeRF-HS is evaluated using peak signal-to-noise ratio (PSNR), structural similarity index measure (SSIM), and root mean squared error (RMSE). PSNR and SSIM measure the spatial reconstruction quality of the rendered views, whereas RMSE assess the spectral accuracy of the reconstructed HSI. All methods are evaluated using the same train test split.

To demonstrate the effectiveness of the proposed BoNeRF-HS framework, we compare it with relevant NeRF based models. NeRF–^25^ model is extended from RGB to *B*-band hyperspectral radiance prediction. SiNeRF^28^ is included as a sinusoidal neural-field for joint pose and radiance-field optimisation. NoPe-NeRF^29^ is also used as a comparison to learn a radiance field without externally provided camera poses. For all comparison methods, the RGB radiance output is replaced by a *B*-band hyperspectral output to enable direct comparison on the same hyperspectral reconstruction task. Table 1 reports the reconstruction results for the mouse bone, trabecular bone analogue, and cortical bone analogue samples. BoNeRF-HS achieves the best PSNR and SSIM on all three samples, indicating improved spatial reconstruction quality compared with NeRF–^25^, SiNeRF^28^, and NoPe-NeRF^29^. On the mouse bone sample, BoneHS-NeRF obtains a PSNR of 29.94, an SSIM of 0.9254, and the lowest RMSE of 0.0325. On the trabecular bone analogue, it achieves the highest PSNR and SSIM, while matching the best RMSE of 0.0547. On the cortical bone analogue, BoneHS-NeRF again gives the best performance, with a PSNR of 31.78, an SSIM of 0.9232, and the lowest RMSE of 0.0258. These results show that the proposed framework improves hyperspectral reconstruction across both real bone and bone analogue samples.

**Table 1.** Quantitative comparison across the three hyperspectral samples. Higher PSNR and SSIM indicate better reconstruction quality, while lower RMSE indicate better spectral accuracy.

| Method | Mouse bone |  |  | Trabecular bone analogue |  |  | Cortical bone analogue |  |  |
| --- | --- | --- | --- | --- | --- | --- | --- | --- | --- |
|  | PSNR↑ | SSIM↑ | RMSE↓ | PSNR↑ | SSIM↑ | RMSE↓ | PSNR↑ | SSIM↑ | RMSE↓ |
| NeRF<sup>-25</sup> | 27.87 | 0.8992 | 0.0408 | 23.75 | 0.7540 | 0.065 | 27.34 | 0.8992 | 0.0444 |
| SiNeRF<sup>28</sup> | 22.84 | 0.8439 | 0.0729 | 25.24 | 0.7864 | **0.0547** | 31.23 | 0.9177 | 0.0274 |
| NoPe-NeRF<sup>29</sup> | 25.83 | 0.8518 | 0.0518 | 24.96 | 0.7742 | 0.0564 | 30.94 | 0.9136 | 0.0287 |
| BoNeRF-HS (Ours) | **29.94** | **0.9254** | **0.0325** | **26.13** | **0.7919** | **0.0547** | **31.78** | **0.9232** | **0.0258** |

Figure 1 presents qualitative reconstruction results at spectral band 50 for the mouse bone, trabecular, and cortical bone analogues. BoNeRF-HS is compared with NeRF–^25^, SiNeRF^28^, and NoPe-NeRF^29^. For each sample, the top row shows the rendered HSI band image, while the bottom row shows the absolute difference heatmap with respect to the ground truth. The red boxes mark selected regions of interest (ROI), and the zoomed insets provide enlarged views of these regions to highlight local structural detail, edge sharpness, and texture preservation.

**Figure 1.**
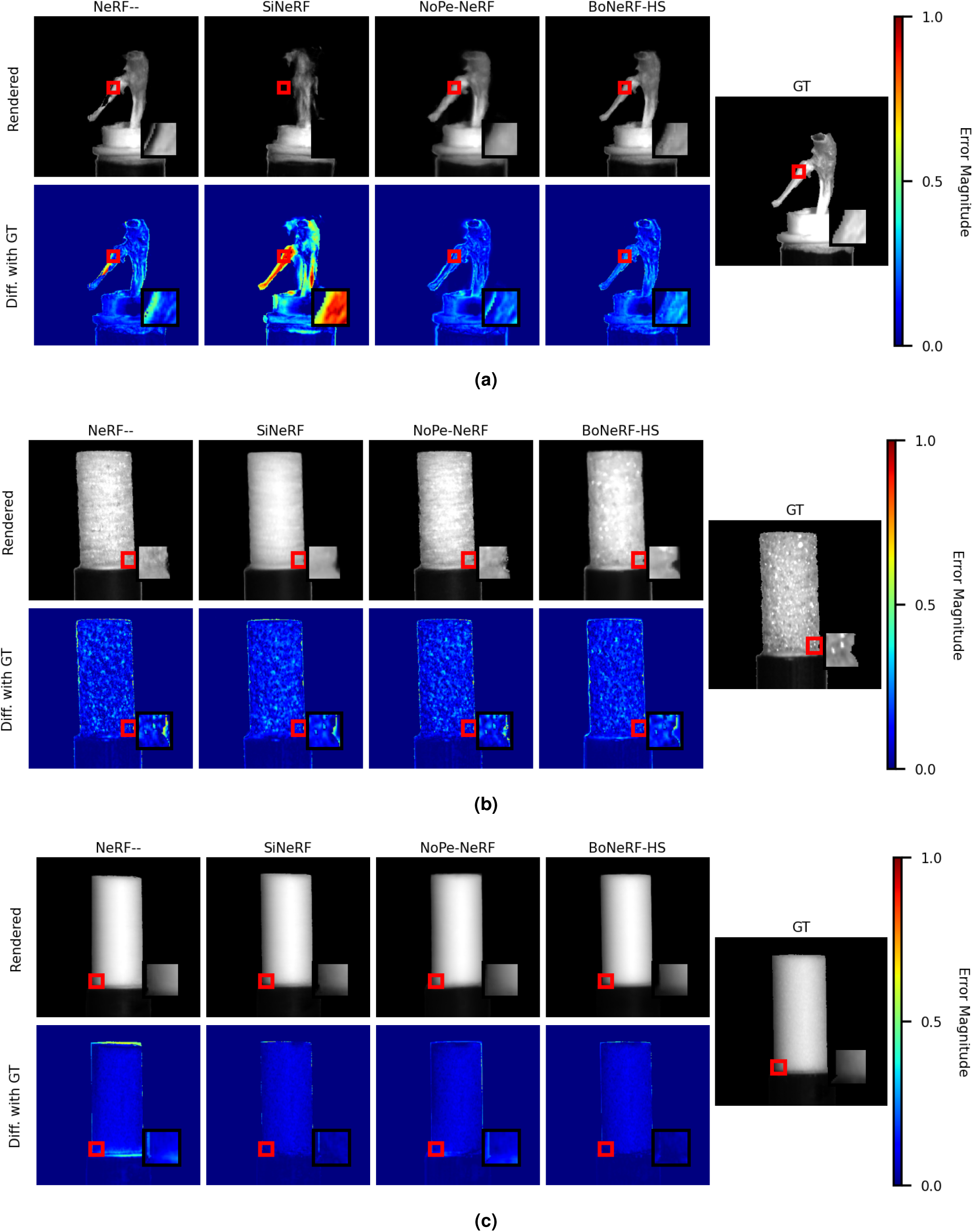
Qualitative comparison of reconstruction results at spectral band 50 for (a) mouse bone, (b) trabecular bone analogue, and (c) cortical bone analogue. The top row shows reconstructed images from NeRF–^25^, SiNeRF^28^, NoPe-NeRF^29^, BoNeRF-HS, and the ground truth. The bottom row shows absolute error maps withrespect to the ground truth, where brighter colours correspond to larger errors. The red boxes indicate selected regions of interest, and the zoomed insets provide enlarged views of these regions to highlight local structural details, edge sharpness, and texture preservation.

For the mouse sample, the compared methods recover the overall object shape, but their error maps show visible residuals around the bone surface, object boundaries, and fine local structures. SiNeRF^28^ produces a smoother appearance but loses some local surface detail, while NeRF–^25^ and NoPe-NeRF^29^ preserve the global shape but still show noticeable differences from the ground truth. In the zoomed ROI, BoNeRF-HS preserves the selected surface region more clearly and produces lower magnitude errors around the highlighted area.

For the trabecular bone analogue, the porous and textured surface makes reconstruction more challenging. NeRF–^25^, SiNeRF^28^, and NoPe-NeRF^29^ reproduce the main cylindrical shape, but the heatmaps show scattered residual errors across the surface texture. The zoomed ROI further shows that local texture and porous surface details are more difficult to preserve. BoNeRF-HS provides a more consistent reconstruction in the highlighted region and reduces the magnitude of the residual errors, indicating improved preservation of local spectral and structural detail.

For the cortical bone analogue, all methods reconstruct the compact cylindrical shape. However, the difference heatmaps reveal residual errors along the object boundary and surface edges. NeRF–^25^ shows stronger boundary artefacts, while SiNeRF^28^ and NoPe-NeRF^29^ still retain visible local errors. BoNeRF-HS produces a cleaner reconstruction with lower visible error, in the highlighted ROI and along the compact bone boundary. Overall, the rendered images, heatmaps, and zoomed regions show that BoNeRF-HS better preserves fine surface details and produces more spatially consistent hyperspectral reconstructions across the three samples.

Figure 1 presents the rendered HSI and absolute difference heatmaps for mouse bone, trabecular bone, and cortical bone samples. BoneHS-NeRF is compared with NeRF–^25^, SiNeRF^28^, and NoPe-NeRF^29^. For each sample, the rendered images show the reconstructed appearance at the selected spectral band, while the difference heatmaps show the absolute error with respect to the ground truth. For the rat bone sample, NeRF–^25^, SiNeRF^28^, and NoPe-NeRF^29^ recover the overall object shape, but their error maps show visible residuals around the bone surface, object boundaries, and fine anatomical structures. SiNeRF^28^ produces a smoother reconstruction but loses some local surface detail, while NeRF–^25^ and NoPe-NeRF^29^ preserve the global structure but still exhibit noticeable differences from the ground truth. In comparison, BoNeRF-HS produces a sharper rendering that is visually closer to the ground truth and shows lower magnitude errors around the fine surface boundaries.

For the trabecular bone analogue, the porous and textured structure makes reconstruction more challenging. NeRF–^25^, SiNeRF^28^, and NoPe-NeRF^29^ reproduce the main cylindrical shape, but the heatmaps reveal scattered errors across the surface texture and internal porous regions. BoNeRF-HS provides a more consistent reconstruction of the trabecular bone analogue and reduces the magnitude of the residual errors, indicating improved preservation of local spectral and structural detail.

For the cortical bone analogue, all compared methods recover the compact cylindrical geometry. However, the difference heatmaps show that NeRF–^25^ produces stronger boundary errors, while SiNeRF^28^ and NoPe-NeRF^29^ still retain visible residuals along the surface. BoNeRF-HS produces the lowest visible error, particularly around the surface and boundary regions. This suggests that the proposed method better preserves the dense compact bone structure.

Figure 2 evaluates the spectral reconstruction quality of the compared methods across selected hyperspectral bands and pixel-wise spectral curves. NeRF–^25^, SiNeRF^28^, and NoPe-NeRF^29^ reproduce the general appearance of the samples, but their spectral curves show visible deviations from the ground truth, indicating spectral inconsistency across wavelength bands. BoNeRF-HS produces spectral curves that more closely follow the ground truth.

**Figure 2.**
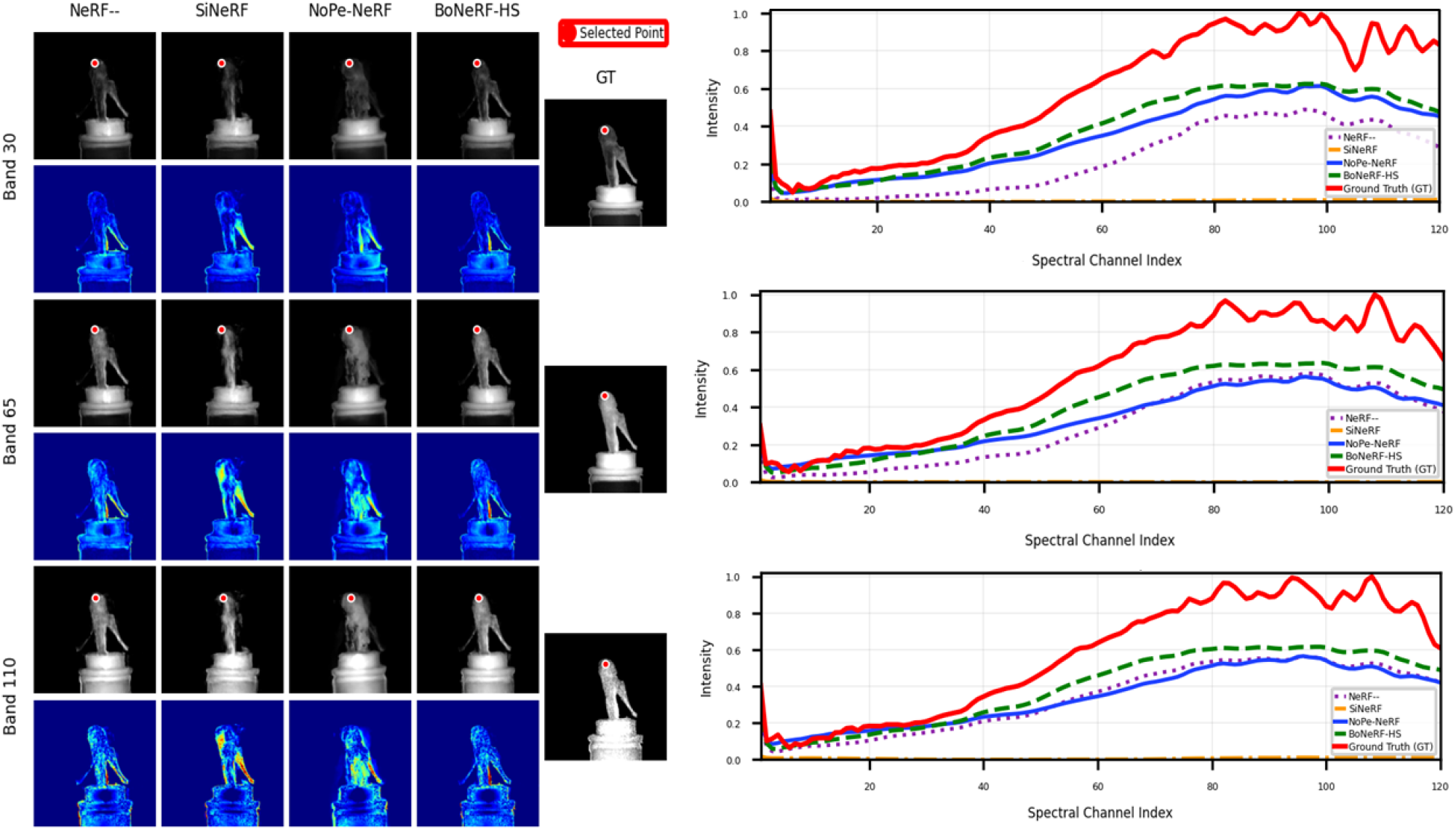
Qualitative results on the mouse knee scene across three different bands 30, 65 and 110. The rendered images and difference heatmaps against the ground truth demonstrate the spectral fidelity and spatial consistency of the reconstruction results, particularly under challenging near-infrared and ultraviolet conditions. In addition, we visualize the reconstructed pixel intensities across all the spectral channels of three randomly selected points in the rightmost column. Compared to the baselines, our method exhibits the highest similarity to the ground truth.

## Discussion

In this section, we explore further experiments and discussions on the impact of gated spectral adapter head and the application of the proposed model in soft tissue reconstruction.

### Impact of the Gated Spectral Adapter Head

To analyse the effect of the gated spectral adapter head, we compare the direct spectral head with three adapter based variants: adapter without gate, adapter with fixed weak gate, and adapter with learned gate as shown in Table 2. This ablation evaluates whether residual spectral correction improves the spectral prediction and whether gating is useful for controlling the magnitude of the residual correction. The adapter design ablation shows that residual spectral correction improves the direct spectral head when it is properly controlled. The fixed weak gate performs worse than the other adapter variants, suggesting that a single constant correction strength is less flexible than an adaptive gate. These results indicate that the spectral adapter is beneficial, but the magnitude of its residual correction must be carefully controlled.

**Table 2.** Ablation study of the spectral adapter design in BoneHS-NeRF, averaged over mouse bone, trabecular and cortical bone analogues. DSH denotes the direct spectral head, SA denotes the spectral adapter, and Gate denotes whether the residual spectral correction is controlled before being added to the direct spectrum. Best results are shown in bold.

| Method | DSH | SA | Gate | PSNR ↑ | SSIM ↑ | RMSE ↓ |
| --- | --- | --- | --- | --- | --- | --- |
| Direct spectral head | ✓ | ✗ | ✗ | 26.32 | 0.8428 | 0.0500 |
| Adapter without gate | ✓ | ✓ | ✗ | 27.37 | 0.8549 | 0.0439 |
| Adapter with fixed weak gate | ✓ | ✓ | ✓ | 25.90 | 0.8190 | 0.0564 |
| Adapter with learned gate (Ours) | ✓ | ✓ | ✓ | **29.29** | **0.8810** | **0.0376** |

### Performance Analysis

Additional performance analysis was conducted on a mouse tissue sample captured using the same data acquisition and preprocessing procedures described in dataset Acquisition and methods Section. For comparison, BoNeRF-HS was evaluated against NeRF–^25^, SiNeRF^28^, and NoPe-NeRF^29^, as reported in Table 3. The proposed BoNeRF-HS model achieved the best overall performance and produced more consistent reconstruction results than the existing approaches. Figure 3 further shows the rendered HSI bands and absolute difference heatmaps for the mouse tissue sample. The compared methods recover the general appearance of the tissue, but their error maps show visible residuals near the boundaries of the tissue, textured regions, and fine local structures. In contrast, BoNeRF-HS produces a sharper rendering that is visually closer to the ground truth and shows lower magnitude errors.

**Table 3.**
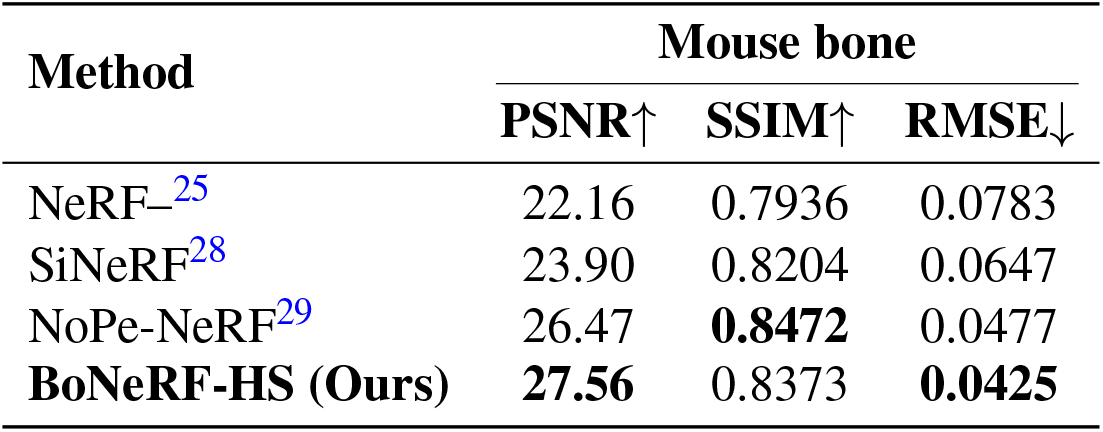
Quantitative comparison on the mouse tissue sample. Higher PSNR and SSIM indicate better reconstruction quality, while lower RMSE indicates better intensity accuracy.

**Figure 3.**
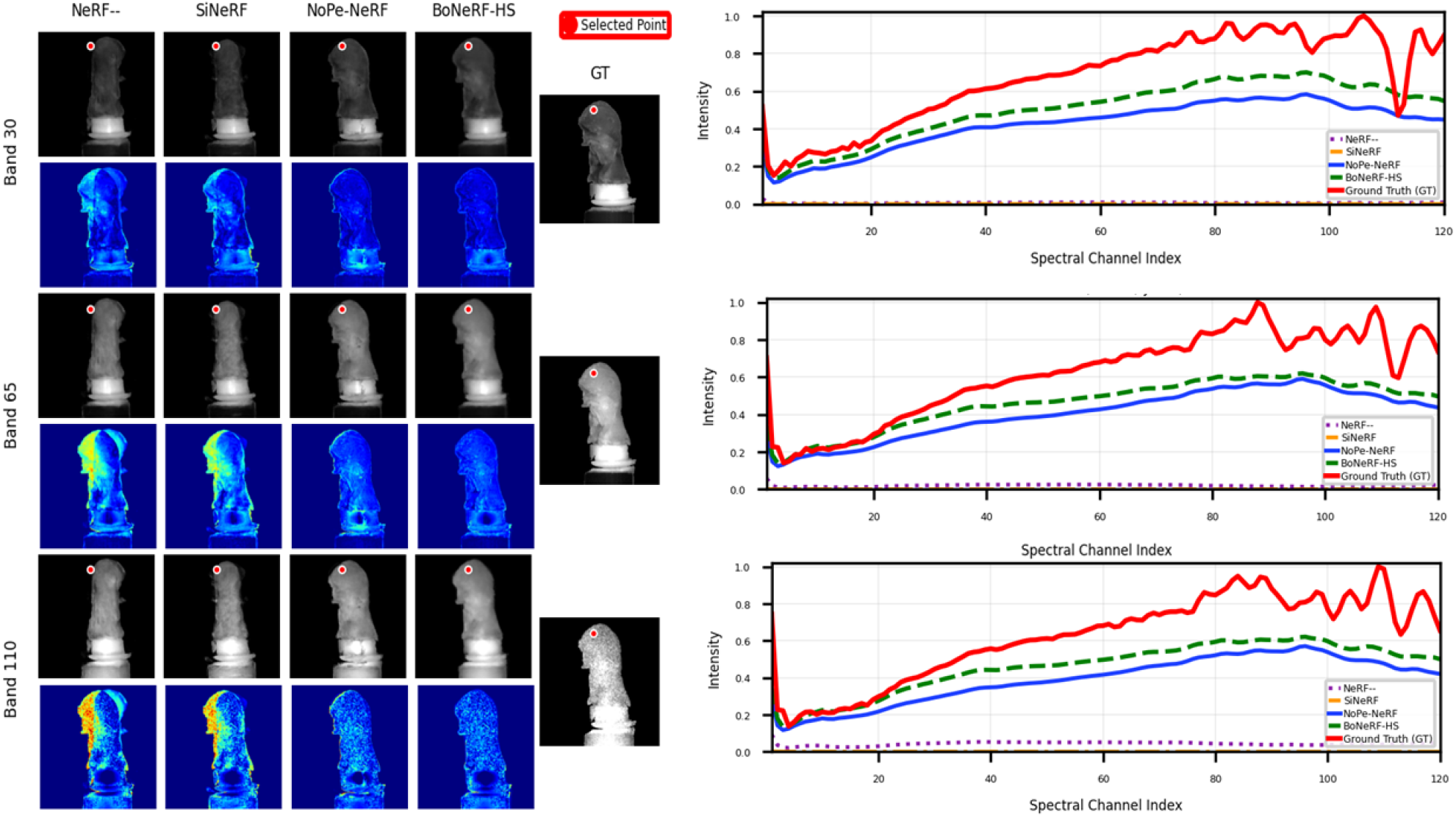
Qualitative results on the mouse tissue scene across three different bands 30, 65 and 110. Each column shows renderings and difference maps for NeRF–^25^, SiNeRF^28^, NoPe-NeRF^29^ and proposed method. n addition, we visualize the reconstructed pixel intensities across all the spectral channels of three randomly selected points in the rightmost column. Compared to the baselines, our method exhibits the highest similarity to the ground truth.

HSI captures rich wavelength dependent information, which is important for analysing mineralised tissues. In this work, we propose BoNeRF-HS, a self-calibrated hyperspectral neural radiance field for 3D bone reconstruction. In conclusion, most current HSI 3D reconstruction methods rely on externally estimated camera poses or RGB based radiance modelling, which may limit their applicability to multi view hyperspectral acquisition. In contrast, BoNeRF-HS jointly optimises camera intrinsics, volume density, and hyperspectral radiance, allowing accurate reconstruction without COLMAP poses. To better utilise the spectral information present in HSI data, we incorporate a gated spectral adapter head that learns wavelength dependent residual information while preserving the direct spectral prediction. We demonstrated the effectiveness of BoNeRF-HS using multi-view HSI data from mouse bone, trabecular, and cortical bone analogues. The results show that the proposed framework improves hyperspectral reconstruction quality compared with the existing methods, producing sharper rendered views and better preservation of fine bone surface details.

Beyond the reconstruction experiments, BoNeRF-HS may have broader relevance in clinical, forensic, and biomechanical applications. Clinically, the framework could support non-contact 3D spectral visualisation of exposed bone surfaces, where both surface structure and tissue specific spectral response are important for future image guided orthopaedic applications^10,11,17,19^. Since our experiments include mouse tissue, mouse bone, trabecular, and cortical bone analogues, the results provide an initial step towards evaluating self-calibrated 3D hyperspectral reconstruction across both biological tissue and controlled bone like materials.

The framework may also be useful in forensic science, where spectral and shape based analysis of skeletal material has been explored for post-mortem interval estimation^30^, and human remain classification^31,32^. In this context, BoNeRF-HS could provide a non-contact approach for documenting skeletal samples while preserving both surface morphology and wavelength dependent spectral information. Further validation on larger and more diverse tissue, bone, and forensic datasets is required before translation to these applications.

## Dataset Acquisition and Methods

In this section, we introduce the hyperspectral dataset used in this study. We then present the proposed BoNeRF-HS framework, a self-calibrated hyperspectral neural radiance field designed for 3D reconstruction.

### Dataset and Preprocessing

We acquire multi-view hyperspectral images using a hyperspectral camera and a turntable motorized rotational sample stage. All hyperspectral data were acquired using a NIREOS HERA VNIR hyperspectral camera^33^. The camera operates in the visible and near-infrared (NIR) of 400 −1000 nm. In this work, each HSI was exported with 120 spectral bands and spatial resolution of 320 × 332 pixels.

The dataset consists of three specimen types, one mouse bone and two bone analogues, namely a trabecular and a cortical bone analogues. Mouse knee were dissected following euthanisation via an appropriate Schedule 1 technique (as listed by the UK Home Office). The tibiofemoral knee joints were then fixed for 48 hours in 4% paraformaldehyde (PFA) and washed in phosphate buffered saline, prior to being trimmed to remove all the exterior muscle and expose the underlying bone for imaging. The bone analogues were prepared from Sawbones polyurethane foams (Pacific Research Laboratories, WA, USA), using open-cell PCF15 foam to represent trabecular-like porous bone and closed cell PCF40 foam to represent denser cortical-like bone. Cylindrical samples of 8 mm diameter and 10 mm length were milled from the foam blocks. Trabecular bone is characterised by a porous cancellous architecture formed by interconnected struts and rods, whereas cortical bone is a denser compact structure with much lower porosity^34,35^. The use of bone analogue materials is motivated by previous phantom and biomaterial studies, where synthetic or cast materials are designed to approximate selected physical or imaging properties of bone^36, 37^.

For data acquisition, mouse bone and bone analogue specimens were mounted on a sample holder and placed on the motorized rotational sample stage. The camera is kept fixed under constant illumination, while the object is rotated to obtain HS images from multiple viewing directions. Each specimen contains 45 hyperspectral views acquired from 0^◦^ to 360^◦^ with approximately 8^◦^ angular spacing.

The acquired HSI is preprocessed. For reflectance calibration, the raw hyperspectral image *I*_raw_ was corrected using a dark reference *I*_dark_ and a white reference *I*_white_. The dark reference was captured with the camera lens closed and the illumination switched off, while the white reference was captured using a reference white target. The calibrated reflectance image *I*_ref_ was computed as

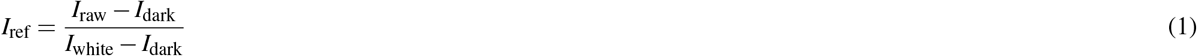

Background removal was applied during preprocessing. Since the camera and illumination were fixed and only the specimen rotated, the background remained stationary with respect to the camera and did not rotate with the object. This could introduce inconsistent background information across views. Therefore, a foreground mask was generated for each hyperspectral view and applied uniformly to all spectral bands. Pixels outside the foreground region were set to zero, producing a pure black background in every wavelength channel. The final processed dataset for each specimen contains 45 calibrated HS views.

### Methods

Figure 4 illustrates the proposed BoNeRF-HS framework. The input to the proposed framework is a set of multi-view HS image captured from different viewpoints of the same specimen. Let **I**_*i*_ ∈ ℝ^*H*×*W* ×*B*^ denote the *i*-th hyperspectral image cube in the acquired multi-view dataset, where *H* and *W* denote the image height and width, respectively, and *B* denotes the number of spectral bands. The model is trained using the HSI views directly, rather than converting them to RGB images. This allows the network to model the wavelength dependent appearance across all *B* spectral bands. Each component of proposed framework describe in this section.

**Figure 4.**
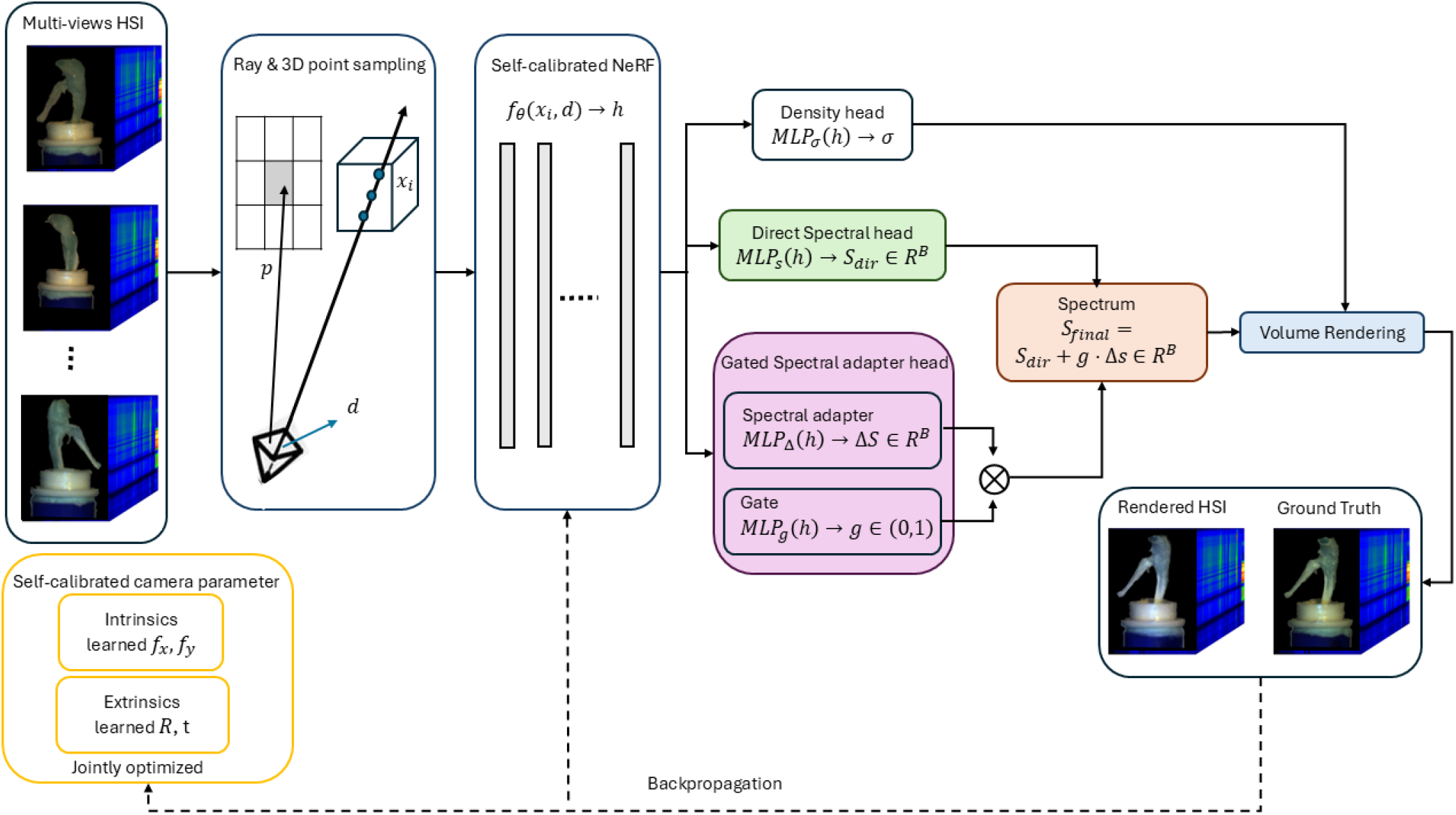
Proposed BoNeRF-HS pipeline for self-calibrated hyperspectral 3D reconstruction. Multi-view HS images with spectral bands *B* are used as input. Camera intrinsics and extrinsics are jointly optimized with the NeRF model, avoiding the need for externally provided camera poses. The NeRF backbone (vertical bar denote fully connected MLP layers) maps sampled 3D points and viewing directions to a latent feature *h*, from which a density head predicts *σ*, a direct spectral head predicts *S*_dir_, and a gated spectral adapter predicts Δ*S* and *g*. The spectrum is *S*_final_ = *S*_dir_ + *g*.Δ*S*, which is volume-rendered to reconstruct the HS images.

### Self-Calibrated Camera Parameter

Conventional NeRF method assumes known camera intrinsics and extrinsics^20,38^. In contrast, BoNeRF-HS framework does not require externally estimated camera parameters. Inspired by NeRF−− ^25^, the proposed BoNeRF-HS framework avoids the need for externally provided camera calibration. The camera intrinsics and per view extrinsics are treated as learnable parameters and are optimised together with the hyperspectral radiance field.

For each view *i*, the camera intrinsics are parameterised by learnable focal lengths *f*_*x*_ and *f*_*y*_. The intrinsic matrix is defined as

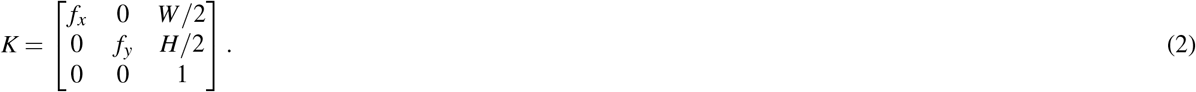

The extrinsic parameters of view *i* are represented by a camera-to-world transformation

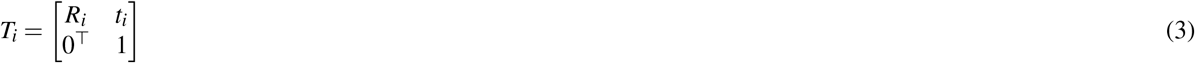

where *R*_*i*_ is the camera rotation and *t*_*i*_ is the camera translation vector.

Both the camera intrinsics (*f*_*x*_, *f*_*y*_) and the extrinsics (*R*_*i*_, *t*_*i*_) are optimised during training. Therefore, the model does not require externally estimated camera calibration or camera poses.

### Ray Generation and 3D Point Sampling

For a pixel location *p* in image *i*, a camera ray *r* is generated using the current estimates of the camera intrinsics and extrinsics. The ray is defined as

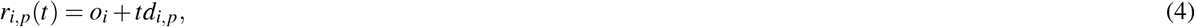

where *f o*_*i*_ is the camera centre, *d*_*i,p*_ is the viewing direction passing through pixel *p*, and *t* denotes the distance along the ray. A set of *N*_*s*_ sample points is drawn along each ray

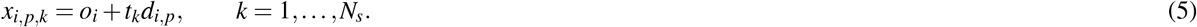

Each sampled point *x*_*i,p,k*_ and the viewing direction *d*_*i,p*_ are used as inputs to the hyperspectral NeRF model.

### Self-Calibrated Hyperspectral NeRF

The NeRF model is implemented as a multilayer perceptron (MLP)^39^, following the standard NeRF model^20^. The sampled 3D point and viewing direction are first encoded using positional encoding,

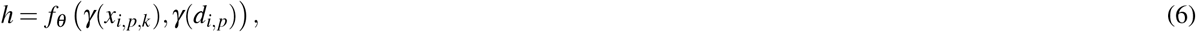

where *γ*(·) denotes positional encoding, *f*_*θ*_ is the NeRF backbone, and *h* is the latent feature vector. This shared feature is passed to three prediction heads: a density head, a direct spectral head, and the proposed gated spectral adapter head.

### Density Head

The density head predicts the volume density at each sampled 3D point,

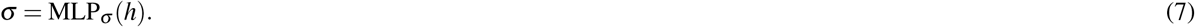

The density *σ* controls how much each sampled point contributes to the final rendered ray. This follows the volumetric density modelling used in NeRF^20^.

### Direct Spectral Head

The original NeRF^20^ predicts RGB radiance. To support HSI, we replace the RGB colour head with a *B*-band spectral head. This is consistent with hyperspectral NeRF model that extend radiance prediction to multi-channel or wavelength dependent spectral outputs^21,40^. The direct spectral head maps the latent feature *h* to an initial *B* band spectrum,

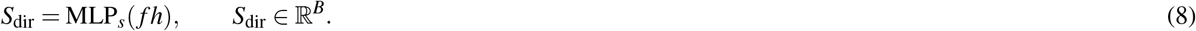

Here, *S*_dir_ denotes the direct hyperspectral radiance predicted by the model before the adapter correction.

### Gated Spectral Adapter Head

Although the direct spectral head predicts a full *B* band hyperspectral spectrum, it treats the spectral bands primarily as output channels. HSI contain many contiguous wavelength bands, and neighbouring bands often exhibit strong spectral correlation rather than being independent. Therefore, modelling the spectral output as a structured wavelength dependent signal is important for preserving material specific spectral signatures^41,42^. To better capture these inter-band dependencies, we introduce a gated spectral adapter head after the NeRF backbone. It is inspired by the general idea of lightweight adapter and low-rank model adaptation^43^.

The spectral adapter predicts a residual correction Δ*S* ∈ ℝ ^*B*^,

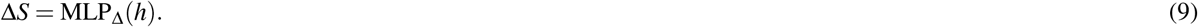

In parallel, a gate branch predicts a confidence value *g* ∈ (0, 1),

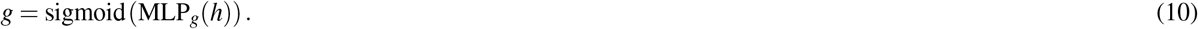

The final spectrum used for rendering is computed as

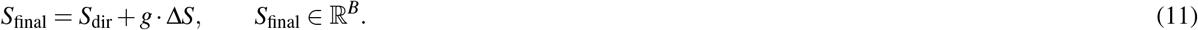

This Eq. 11 allows the adapter to learn a gated wavelength correlated correction.

### Hyperspectral Volume Rendering

For each sampled point along a ray, the model predicts a density value *σ*_*k*_ and a final hyperspectral spectrum *S*_final,*k*_. We render the final hyperspectral pixel using the differentiable volume rendering formulation adopted from NeRF^20^. The opacity of sample *k* is

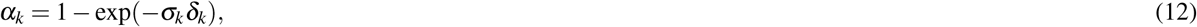

where *δ*_*k*_ is the distance between adjacent sampled points. The accumulated transmittance is

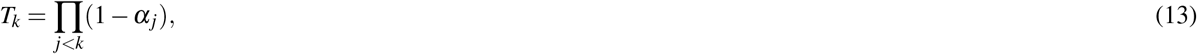

and the rendering weight is

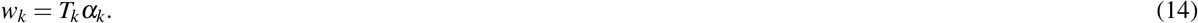

The rendered hyperspectral pixel is obtained by integrating the *B*-band spectra along the ray,

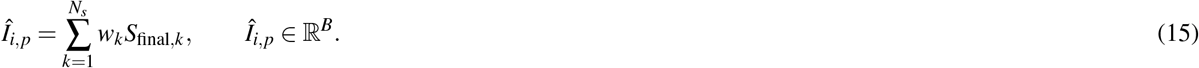

Applying this operation to all pixels produces the rendered hyperspectral image

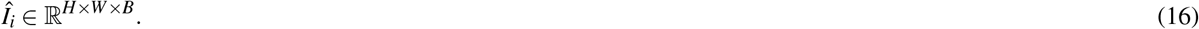

## Data availability

The authors declare that all data supporting the current study are available under reasonable request.

## Acknowledgements

The authors gratefully acknowledge NIREOS for providing HERA hyperspectral camera^33^ used for data collection in this study.

## Ethics declarations

Mouse knee joints from male one year old C57BL/6 were obtained from Queen Mary University of London. All procedures complied with the UK Animals (Scientific Procedures) Act of 1986 and were performed under a UK Home Office Project Licence in accordance with the Queen Mary University of London Ethical Policy on the use of Animals in Scientific Research and the ARRIVE guidelines. The study was approved by ethical review board at the Queen Mary University of London. Mice were housed under standard conditions with a 12 h light/dark cycle, and unlimited access to food and water.

## Author contributions statement

G.T. initialised concepts and directions. S.A. provided the mouse sample. G.T., T.N., and N.S. acquired the dataset. N.S. conducted the experiments and analysed the results. T.N. and G.T. provided critical updates and suggestions that significantly enhanced the scope and direction of the research. All authors N.S., T.N., S.A. and G.T. reviewed and approved the final manuscript.

